# Haplotype-resolved T2T gap-free genomes of the winegrape cultivar ‘Cabernet Sauvignon’

**DOI:** 10.1101/2025.07.26.666927

**Authors:** Falak Sher Khan, Tiepeng Sun, Xiangfeng Wang, Xuenan Zhang, Naomi Abe-Kanoh, Ying Hua Su, Li Guo, Wenxiu Ye

**Affiliations:** State Key Laboratory of Wheat Improvement, College of Life Sciences, Shandong Agricultural University, Tai’an, 271018, Shandong, China; Shandong Key Laboratory of Precision Molecular Crop Design and Breeding, Peking University Institute of Advanced Agricultural Sciences, Shandong Laboratory of Advanced Agricultural Sciences in Weifang, Weifang, 261325, China; United Graduate School of Agricultural Science, Iwate University, 3-18-8 Ueda, Morioka, Iwate, 020-8550, Japan; Department of Food, Life and Environmental Science, Faculty of Agriculture, Yamagata University, Wakaba-machi 1-23, Tsuruoka, Yamagata, 997-8555, Japan

**Author notes:** These authors contributed equally: Falak Sher Khan, Tiepeng Sun.

## Abstract

Cabernet Sauvignon (CS), a cultivar of winegrape (*Vitis vinifera*), is among the most renowned winegrape varieties globally. In this study, we released the haplotype-resolved telomere-to-telomere (T2T) CS genome assembled using a combination of PacBio HiFi, ONT ultra-long, and Hi-C sequencing data. The two T2T gap-free haplotype-resolved assemblies CS-T2T.Hap1 and CS-T2T.Hap2 sized 491.11 Mb (contig N50 = 25.09 Mb) and 491.90 Mb (contig N50 = 25.51 Mb), respectively. Genome annotation predicted a total of 36,456 genes in CS-T2T.Hap1 and 35,471 genes in CS-T2T.Hap2. By genome comparison, we discovered and validated megabase inversion events on Chromosome 03,11,18 and 19, which are not prevalent in other haplotype-resolved *V. vinifera* genomes. In summary, this haplotype-resolved T2T genome represents an essential genomic resource for Cabernet Sauvignon, and lays the foundation for its genetic studies, improvement and utilization.

## Background & Summary

The cultivated grapevine (*V. vinifera* L.) is one of the most valuable perennial fruit crops cultivated for wine-making and fresh consumption worldwide^1^. ‘Cabernet Sauvignon’ is a major red wine cultivar renowned for its thick, resilient skins and high concentration of phenolic compounds. Wine from this variety exhibits a robust tannic structure and complex flavor profile dominated by notes of blackcurrant, plum, and green bell pepper^2^. These traits are attributed to the grapevine’s unique genetic lineage and environmental adaptability, making it one of the most widely cultivated wine grape varieties globally. Originating from a natural cross between Sauvignon Blanc and Cabernet Franc in the Aquitaine region of France prior to the seventeenth century, its historical and viticultural significance is further underscored by its genetic stability and widespread adoption across diverse terroirs^3^.

Since the first *V. vinifera* genome was assembled in 2007 using the grapevine PN40024^4^, originally derived from 9 selfings of ‘Helfensteiner’ (cross of ‘Pinot noir’ and ‘Schiava grossa’)^5^, the field of grapevine genomics has progressed rapidly, with high-quality reference assemblies now available for diverse species and cultivars^6-9^. In 2023, of particular significance, telomere-to-telomere (T2T) genome has been completed for PN40024^10^. Since then, several economically vital varieties such as the white wine grape ‘Chardonnay’^8^, ‘Thomson seedless’^11^, ‘Baimunage’, ‘Black Monukka’, ‘Muscat Hamburg’, ‘Hongmunage’, ‘Manicure Finger’, ‘Thompson’^9^. The genome of ‘Cabernet Sauvignon’ was initially assembled in 2016^12^, followed by the release of several chromosomal-level versions^8,13^.Despite its global prominence as the most widely planted wine grape, ‘Cabernet Sauvignon’ lacks a T2T gap-free reference genome, which hinders comprehensive understanding of the molecular mechanism behind its complex traits, including disease resistance, berry development, and terroir-specific flavor profiles, thus its improvement and utilization.

In this study, we assembled the haplotype-resolved T2T genome of ‘Cabernet Sauvignon’ using PacBio-HiFi, ONT ultra-long, and Hi-C sequencing data. The two haplotype-resolved T2T genome assemblies are gapless and sized 491.11 Mb and 491.90 Mb with N50 length of 25.09 Mb and 25.51 Mb, respectively. A total of 36,456 and 35,471 protein-coding genes were annotated in the two haplotype genomes combining ab initio prediction, homology proteins and transcriptome evidence. Vigorous quality assessment demonstrated the high accuracy and completeness of the assembly and annotation. These T2T assemblies and annotations improve the genomic resources available for *V. vinifera* cultivar ‘Cabernet Sauvignon’ and provide a crucial reference genome for its molecular studies and improvement.

## Methods

### Plant resources and genome sequencing

The ‘Cabernet Sauvignon’ was cultivated in the vineyards at Peking University Institute of Advanced Agricultural Sciences, China. Genomic DNA was extracted from grapevine leaves using the CTAB (cetyltrimethylammonium bromide) method^14^. The Agilent 4200 Bioanalyzer (Agilent Technologies) was used to evaluate the DNA integrity. For HiFi sequencing library preparation, 15 µg of high-molecular-weight genomic DNA was utilized to generate a PacBio SMRTbell library with the SMRT Express Template Prep Kit 2.0 (Pacific Biosciences).Circular consensus sequencing (CCS) was conducted on a PacBio Sequel IIe system (Biomarker Technologies Corporation, Qingdao, China) to yield 87.86 Gb of HiFi data, corresponding to 178 × genome coverage, with read N50 length of 18.06 kb (Table 1). Ultra-long-read sequencing was performed using the Oxford Nanopore Technologies (ONT) platform, where size-selected long DNA fragments were prepared according to the SQK-LSK109 Kit protocol. Sequencing was carried out on the PromethION sequencer at Single-Molecule Sequencing Platform of Annoroad. A total of 49.71 Gb ONT data was generated, corresponding to 101 × genome coverage (Table 1). The Next-Generation Sequencing (NGS) and High-throughput chromosome conformation capture sequencing (Hi-C) datasets used in this study were obtained from published sources, corresponding to 50 × and 126 × genome coverage, respectively^8^ (Table 1).

**Table 1:** Statistic of sequencing data for ‘Cabernet sauvignon’ genome assembly. erage was calculated based on the Hap1 genome assembly of 491.10 Mb.

| Library type | Reads number | Total length (Gb) | N50 length (bp) | Coverage* |
| --- | --- | --- | --- | --- |
| NGS | 165,785,460 | 24.74 | 2 × 150 | 50 × |
| PacBio HiFi | 4,752,646 | 87.86 | 18,062 | 178 × |
| Hi-C | 413,992,094 | 61.78 | 2 × 150 | 126 × |
| ONT | 531,173 | 49.71 | 100,000 | 101 × |

### Genome assembly

To estimate fundamental characteristics of the ‘Cabernet Sauvignon’ genome, such as genome size, heterozygosity, we performed the genome survey using the PacBio HiFi data. The K-mer analysis was performed using Jellyfish v2.3.0^15^ with a K-mer size of 21 and used for the genome survey by GenomeScope v1.0^16^. The genome size of ‘Cabernet Sauvignon’ was estimated at 495.32 Mb, with the heterozygosity rate of 1.56% (Fig. 1a). To assemble the haplotype-resolved T2T genomes for ‘Cabernet Sauvignon’, PacBio HiFi (178 ×), ONT ultra-long (101 ×), and Hi-C (126 ×) reads were integrated to produce an initial assembly of two haplotypes using hifiasm v0.24.0^17^ with the parameters ‘--hom-cov --h1 --h2 --ul’. To eliminate mitochondrial and chloroplast sequences from the genome assembly, the genome contigs were aligned to the *V. vinifera* mitochondrial genome (GenBank: FM179380.1) and chloroplast genome (GenBank: DQ424856.1) using minimap2 v2.28^18^ with the parameter ‘--x asm5’. Contigs exhibiting ≥ 50% alignment coverage to plastid sequences were excluded from the assembly. By comparing contigs with all NCBI RefSeq bacterial genomes using blast-2.13.0+ with the parameter “-task megablast”, other common contamination sequences (such as microbial DNAs) were further eliminated. The high-quality haplotype-resolved assemblies were scaffolded into chromosome-scale genomes using Hi-C data with Juicer^19^ and 3d-DNA^20^ pipeline. Manual corrections to the scaffolded pseudochromosomes were performed using Juicebox v1.11.08^21^, guided by Hi-C chromatin interaction patterns observed in contact heatmaps. To fill the gaps in the chromosome-scale assemblies, ONT ultra-long and HiFi reads were mapped to the haplotype-resolved genomes, and the aligned reads were used to fill gaps. The final genome assembly produced two gap-free haplotypes (Fig. 1b), haplotype 1 (CS-T2T.Hap1, 491.11 Mb) and haplotype 2 (CS-T2T.Hap2, 491.90 Mb) (Table 2) with each genome consists of 19 chromosomes (Fig. 1c). The scaffold N50 lengths for CS-T2T.Hap1 and CS-T2T.Hap2 were 25.09 Mb and 25.51 Mb, respectively (Table 2). The telomere sequences were identified using Tandem Repeat Finder (TRF) v4.09.1^22^, and the result file was converted into GFF3 format via TRF2GFF (https://github.com/Adamtaranto/TRF2GFF) to detect the seven-base repeats. In summary, ‘Cabernet Sauvignon’ assembly can be regarded as having achieved telomere-to-telomere and gap-free completeness.

**Table 2:**
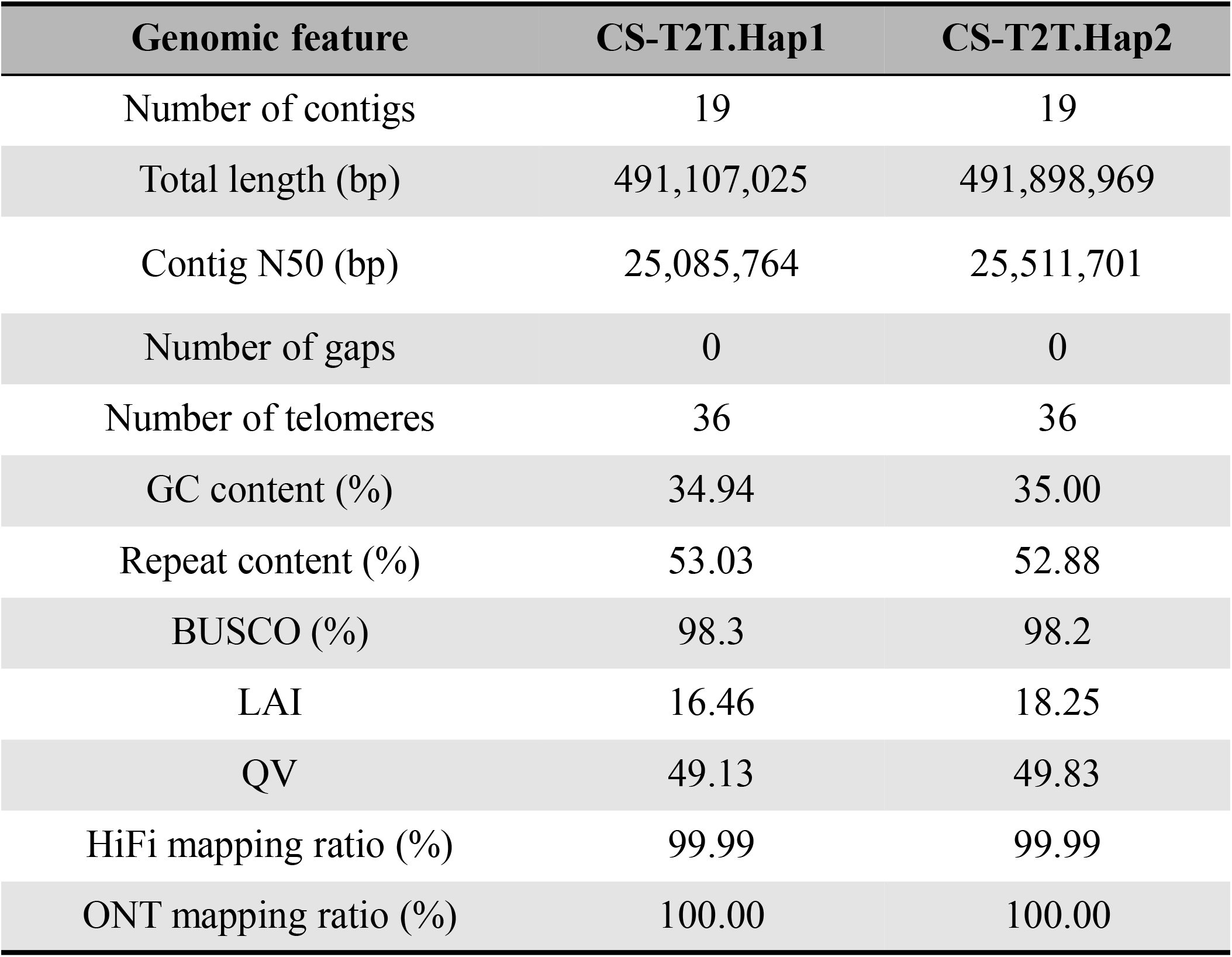
Assembly statistics for the two haplotypes of ‘Cabernet sauvignon’.

| Genomic feature | CS-T2T.Hap1 | CS-T2T.Hap2 |
| --- | --- | --- |
| Number of contigs | 19 | 19 |
| Total length (bp) | 491,107,025 | 491,898,969 |
| Contig N50 (bp) | 25,085,764 | 25,511,701 |
| Number of gaps | 0 | 0 |
| Number of telomeres | 36 | 36 |
| GC content (%) | 34.94 | 35.00 |
| Repeat content (%) | 53.03 | 52.88 |
| BUSCO (%) | 98.3 | 98.2 |
| LAI | 16.46 | 18.25 |
| QV | 49.13 | 49.83 |
| HiFi mapping ratio (%) | 99.99 | 99.99 |
| ONT mapping ratio (%) | 100.00 | 100.00 |

**Figure 1.**
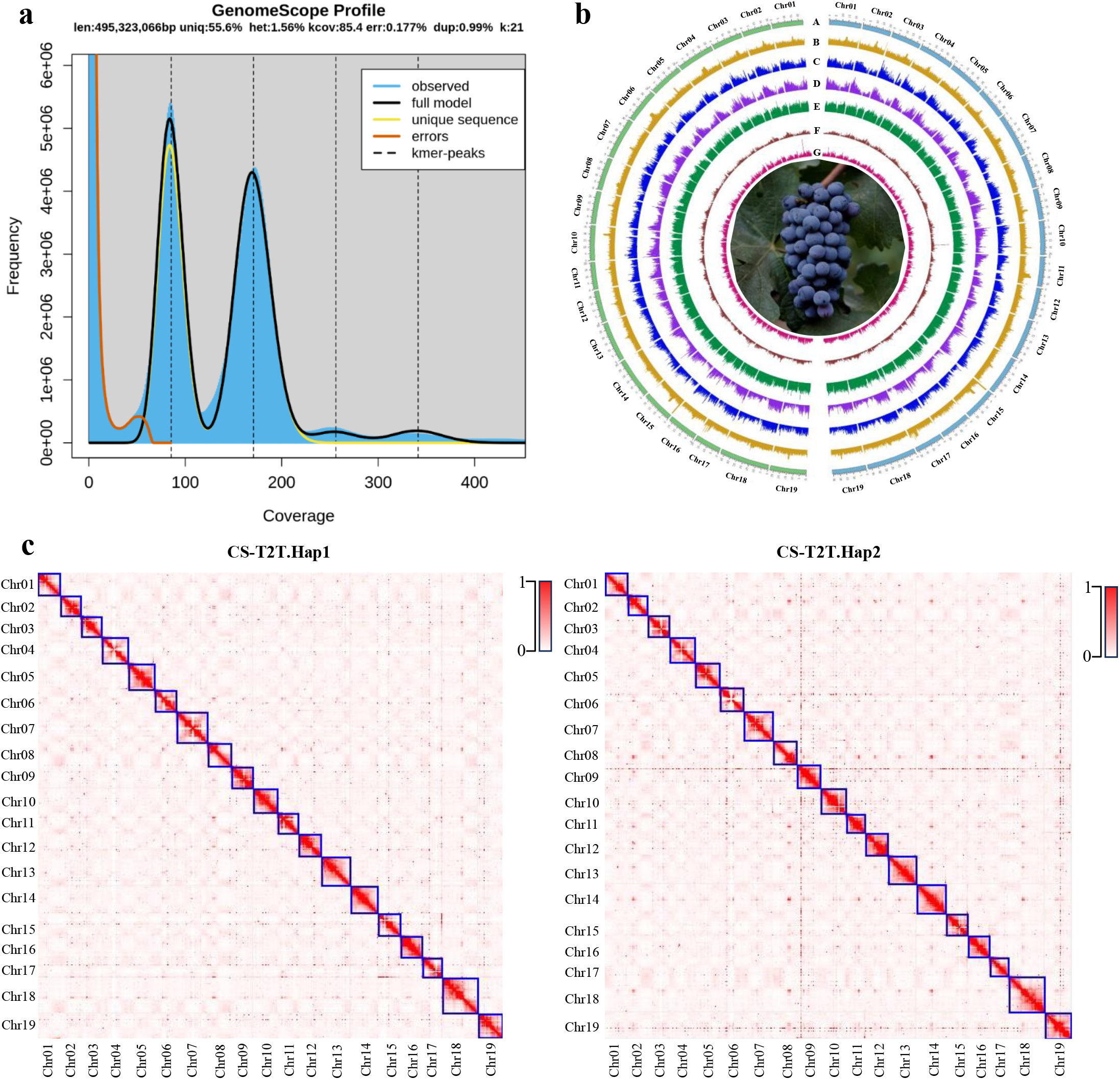
Hapotype-resolved genome assembly of ‘Cabernet Sauvignon’. (a) Genome survey based on K-mer analysis using 1.0. (b) Genomic features of CS genome. Tracks (A) to (G): chromosomes, GC content, gene density, exon density, LTR/Gypsy density, LTR/Copia density. Windows: 50 Kb. Left (green): Haplotype 2. Right (blue):c) Hi-C contact map of CS assembly.

### TE annotation

Before genome annotation, a nonredundant transposable element (TE) library was constructed using the Extensive de novo TE Annotator (EDTA) v2.2.1^23^ with parameters ‘--sensitive 0 -- anno 1 --step all’. For haplotype-resolved assembly, approximately 51.80-52.70% of genome was classified as repetitive elements, which included 39.60-40.60% long terminal repeats (LTRs), 4.07-4.10% long interspersed nuclear elements (LINEs), 0.24-0.25% short interspersed nuclear elements (SINEs), and 7.86-7.81% DNA transposon elements, respectively (Table 3).

**Table 3:**
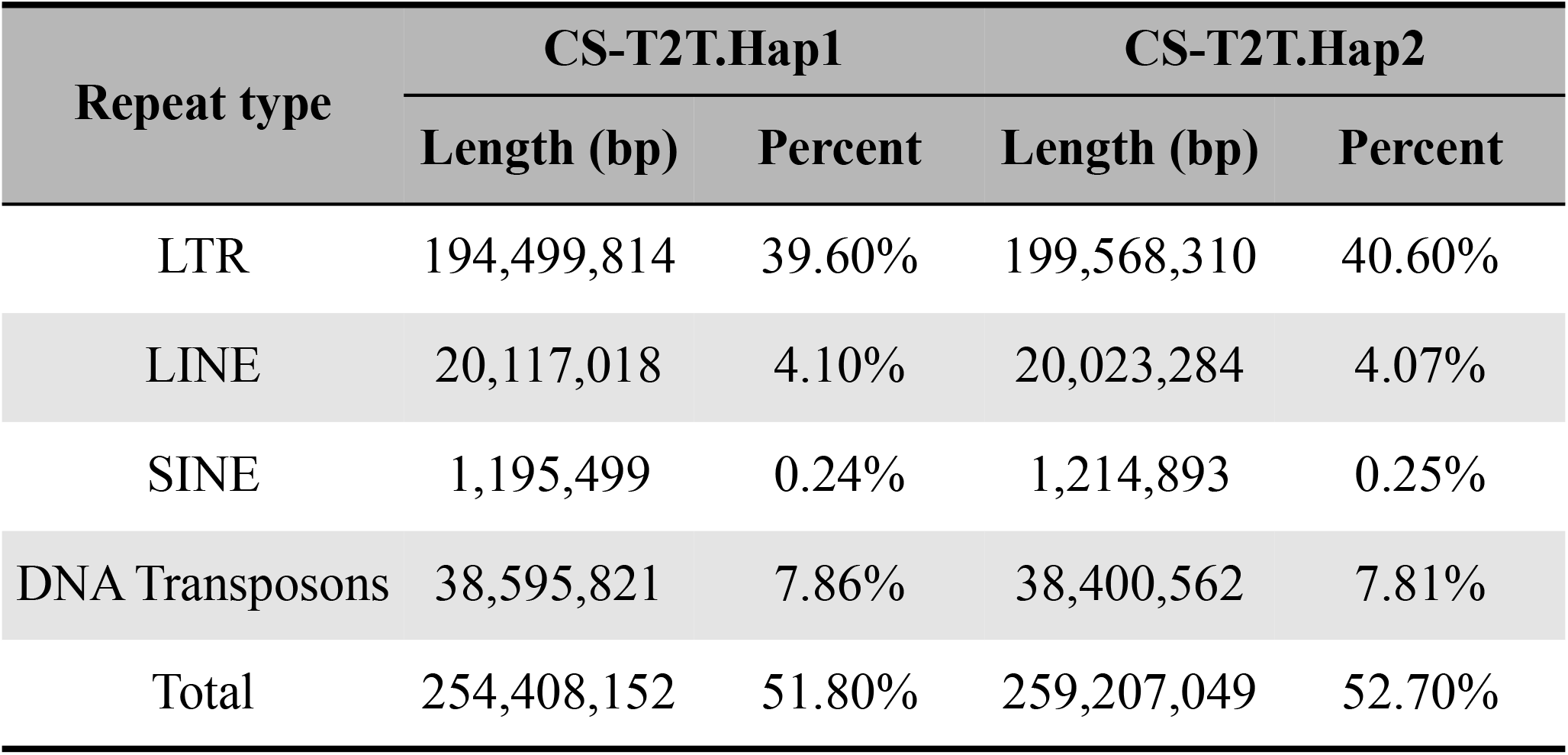
Statistics of annotated repeat elements in the genome of ‘Cabernet Sauvignon‘.

| Repeat type | CS-T2T.Hap1 |  | CS-T2T.Hap2 |  |
| --- | --- | --- | --- | --- |
|  | Length (bp) | Percent | Length (bp) | Percent |
| LTR | 194,499,814 | 39.60% | 199,568,310 | 40.60% |
| LINE | 20,117,018 | 4.10% | 20,023,284 | 4.07% |
| SINE | 1,195,499 | 0.24% | 1,214,893 | 0.25% |
| DNA Transposons | 38,595,821 | 7.86% | 38,400,562 | 7.81% |
| Total | 254,408,152 | 51.80% | 259,207,049 | 52.70% |

### Genome annotation

Gene model prediction was performed using a comprehensive pipeline that integrated multiple data sources after masking repetitive elements, including: ab initio predictions, homologous protein comparisons, and RNA-seq evidence. The ab initio gene prediction was implemented by training the GeneMark-ET model with BRAKER2 v2.1.6^24^ and further refined it by training the semi-HMM model SNAP^25^ via MAKER v3.01.03^26^. The homologous protein sequences were included different published data from *Vitis*^8,10^ and UniProt Swiss-Prot database (https://www.uniprot.org/downloads). The homologous protein dataset was filtered using CD-HIT v4.8.1^27^ with default parameters to remove redundant sequences. The transcriptome data were obtained from published sources and derived from the same cultivar^8^. Transcriptome data were mapped to the haplotype-resolved assembly using HISAT2 v2.2.1^28^, and subsequently processed with StringTie v2.2.1^29^ for genome-guided transcript assembly.The trained gene prediction models, homologous protein sequences and transcriptome evidence were integrated into MAKER to predict gene structures. The final gene model was predicted by the MAKER pipeline with protein-coding genes having an AED < 0.5. A total of 36,456 and 35,471 protein-coding genes were depicted for CS-Hap1 and CS-Hap2, respectively (Table 4).

**Table 4:** Summary of protein-coding genes functional annotation for the two pes of ‘Cabernet sauvignon’.

| Feature | CS-T2T.Hap1 | CS-T2T.Hap2 |
| --- | --- | --- |
| Annotated genes | 36,456 | 35,471 |
| NR | 34,128 | 32,634 |
| eggNOG | 28,549 | 27,110 |
| Pfam | 25,668 | 24,367 |
| Swiss-port | 23,419 | 23,174 |
| KEGG | 13,548 | 13,444 |
| GO | 12,704 | 12,902 |
| Non-redundant annotated | 34,142 | 32,648 |

### Functional annotation of protein-coding genes

The predicted protein-coding genes were functionally annotated utilizing a robust multi-database approach to ensure comprehensive annotation, including Non-Redundant (NR), Evolutionary Genealogy of Genes: Non-supervised Orthologous Groups (eggNOG), Protein Families Database (Pfam), Swiss-Prot protein database, Kyoto Encyclopedia of Genes and Genomes (KEGG) and Gene Ontology (GO) databases. The eggNOG database was engaged via eggNOG-mapper (http://eggnog-mapper.embl.de/). The non-redundant (NR) and Swiss-Prot protein databases were employed for functional annotation using Diamond BLASTP v2.1.6^30^. A total of 34,142 and 32,648 non-redundant protein-coding genes were annotated using the various databases for CS-T2T.Hap1 and CS-T2T.Hap2, respectively (Table 4).

### Genome synteny analysis

To compare published *Vitis* genome^8,10^ with the assembled genome, whole-genome alignments were generated using minimap2 v2.28^18^, followed by sorting and indexing the alignment BAM files with SAMtools v1.20^31^. Genomic synteny and structural variants were identified by SyRI^32^, and the results visualized using Plotsr (https://github.com/schneebergerlab/plotsr). Notably, within the two ‘Cabernet Sauvignon’ haplotype genomes, we discovered megabase-scale inversions, which are not prevalent in other haplotype-resolved *V. vinifera* genomes (Fig. 2a). To validate the chromosomal inversions, we compared the two haplotype genomes of ‘Cabernet Sauvignon’ using minimap2 v2.28^18^ and visualizing by D-genies^33^. For example, a chromosomal inversion of approximately 8 Mb was detected through comparative analysis of chromosome 19 between CS-T2T.Hap1 and CS-T2T.Hap2 (Fig. 2b). Furthermore, chromosomal inversions were validated using Hi-C interaction maps, which confirm a local inversion in this region (Fig. 2b).

**Figure 2.**
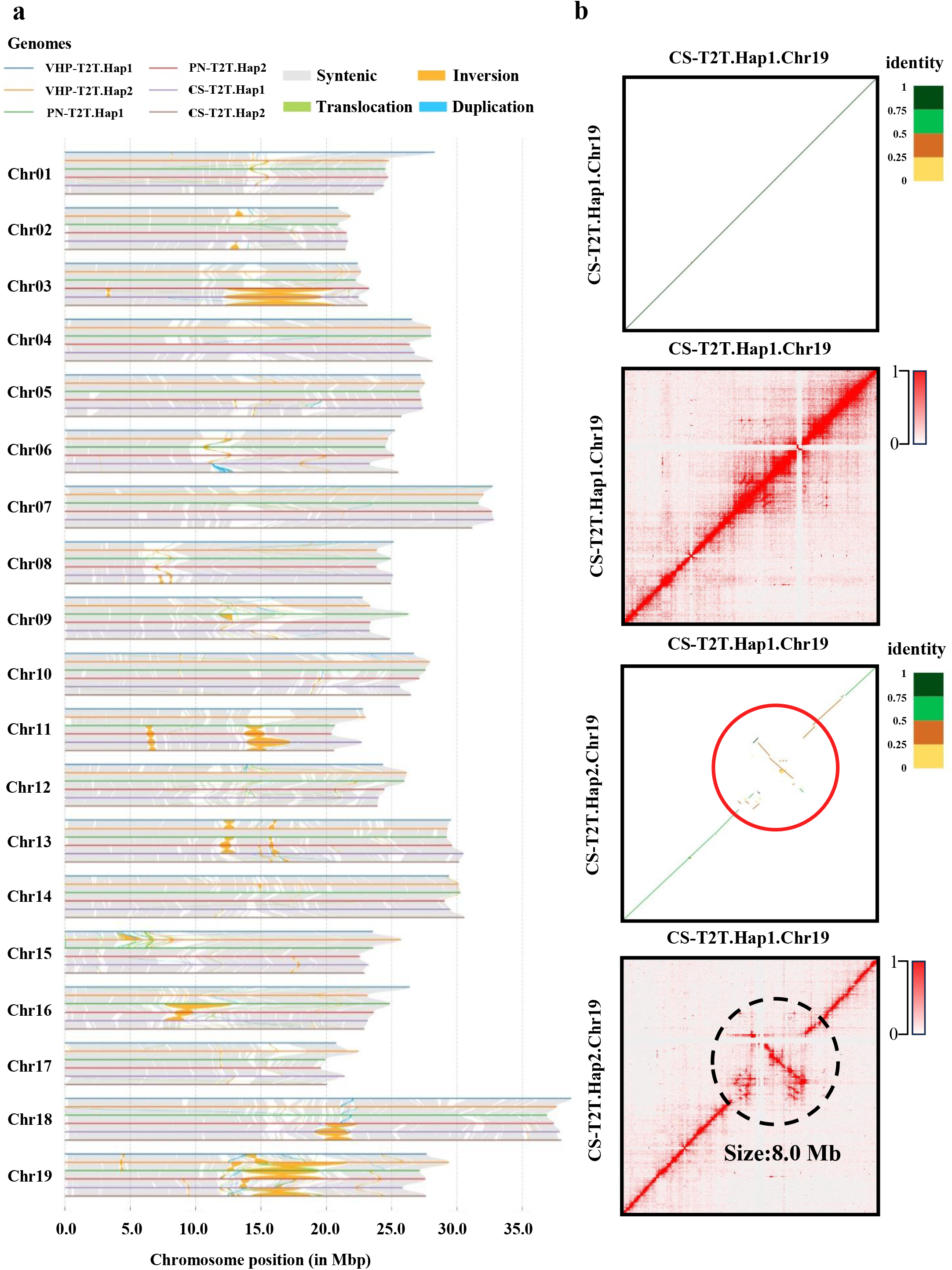
Genome synteny and structural variants analysis. (a) The collinearity and structural variants among six genomes. nay’. PN: ‘PN40024’. CS: ‘Cabernet Sauvignon’. (b) Example of an inversion event on chromosome 19 in two dated by Hi-C data. Up, dot plot showing genome alignment. Down, Hi-C contact heatmap.

### Data Records

The raw genomic sequencing data and genome assemblies have been deposited in the European Molecular Biology Laboratory-European Bioinformatics Institute (EMBL-EBI) (https://www.ebi.ac.uk/) under the study number of PRJEB94081. The accession numbers of PacBio HiFi and ONT sequencing data are publicly accessible as ERR15313518 and ERR15314151 respectively. The accession numbers of genome assemblies are publicly accessible as ERZ27315251 and ERZ27315252 respectively. The annotation files have been deposited in Zenodo (https://doi.org/10.5281/zenodo.16300203).

## Technical Validations

### Genome assembly quality assessment

To evaluate the completeness of the ‘Cabernet Sauvignon’ haplotype-resolved T2T gap-free genome, we employed the Benchmarking Universal Single-Copy Orthologous (BUSCO) (v5.4.2)^34^ with the eudicots_odb10 database. The evaluation of genome completeness demonstrated BUSCO scores of 98.3% and 98.2% for CS-T2T.Hap1 and CS-T2T.Hap2, respectively (Table 2). The ONT and HiFi reads were aligned to genomes with minimap2v2.28 (−ax map-ont, -ax map-pb)^18^. The mapping rate, mean sequencing depth, and genome coverage were calculated via SAMtools v1.20^31^ from BAM files generated by minimap2. The haplotype-resolved genome assemblies showed high mapping ratio: HiFi reads reached 99.99%, and ONT reads achieved 100% mapping to both haplotypes (Table 2), reflecting superior assembly continuity. Long terminal repeats (LTRs) were predicted using LTR_FINDER v1.1^35^, followed by classification and filtering using LTR_retriever v2.9.0^36^ to calculate the LTR Assembly Index (LAI), reflecting genome assembly continuity. LAI scores of 16.46 (CS-T2T.Hap1) and 18.25 (CS-T2T.Hap2) indicated high-quality haplotype-resolved assemblies (Table 2). The k-mer-based quality of the haplotype-resolved assemblies was assessed using Meryl v1.3 and Merqury v1.3^37^. The results revealed quality values (QV) of 49.13 for CS-T2T.Hap1 and 49.83 for CS-T2T.Hap2 (Table 2), indicating high accuracy in both haplotypes. Collectively, these results demonstrate the high accuracy and completeness of the T2T haplotype-resolved genome assembly for the grapevine cultivar ‘Cabernet Sauvignon’.

### Gene annotation assessment

Functional validation of annotated genes was conducted through alignment with different public protein databases. For the two haplotype genomes, revealed that the following proportions of proteins were annotated across databases: NR (92.82%), EggNOG (77.38%), Pfam (69.56%), Swiss-Prot (64.78%), KEGG (37.53%), and GO (35.60%). Overall, 93.65%(Hap1) and 92.04% (Hap2) non-redundant genes were mapped to public protein databases, confirming the comprehensive of the gene annotations.

## Code availability

All software used in this work are publicly available, and the software versions and parameters used are described in the Methods section. The parameters not mentioned in the analysis were used as default parameters suggested by the developer. No specific script was developed in this study.

## Author contributions

W.Y., L.G. and Y.H.S. conceived and supervised the project, X.Z. prepared plant materials.F.S.K. and T.S. performed bioinformatics analysis, prepared the tables, figures, uploaded the raw data and wrote the manuscript. X.W., N.K., L.G. and W.Y. revised the manuscript. All authors have read and approved the final version of the manuscript.

## Acknowledgements

This research was supported by the Key Research and Development Program of Shandong Province (2024LZGC033, 2024CXPT031), and Shandong Provincial Natural Science Foundation (SYS202206), Taishan Scholars Program of Shandong Province and Weifang Key Laboratory of Grapevine Improvement and Utilization, China.

